# Common neural representation between visual perception and imagery in eidetikers

**DOI:** 10.64898/2026.02.18.705196

**Authors:** Ritsuko Nabata, Mitsuki Niida, Kosaku Takahashi, Kenji Ogawa

## Abstract

Eidetic imagery is experienced as “seen” in external space, unlike typical visual mental imagery, which occurs in the mind’s eye. To investigate whether eidetic imagery and perception share common neural substrates, we measured brain activity in eidetikers and typical imagers using fMRI while they performed perception and imagery tasks. Multi-voxel pattern analysis (MVPA) revealed that imagined objects could be classified based on perceptual activation patterns in right BA19 for eidetikers, but not for typical imagers. This effect was specific to BA19 and was not observed in early visual areas (BA17/18). Subjective ratings confirmed that eidetikers experienced images as percept-like. These findings provide neural evidence for percept-like mental imagery and show that individual differences in subjective experience are reflected in the high-level visual cortex.

**Significance Statement:** How does the brain support individual differences in subjective visual experience? Visual mental imagery partially shares neural substrates with perception, but how this shared activity relates to subjective conscious experience remains unclear. Some individuals, known as eidetikers, report images that are literally ‘seen’ and percept-like, providing a unique opportunity to investigate the neural basis of individual variation. Using fMRI and multivariate pattern analysis, we show that in eidetikers, the neural representations of imagined objects are more strongly shared with perceptual representations in a higher-level visual area (BA19) than in typical imagers. These findings provide neural evidence for eidetic imagery and reveal how individual differences in subjective experience are reflected in the organization of visual representations in the human brain.

## Introduction

Understanding how neural activity gives rise to variability in subjective experience is a central goal of neuroscience. Early work by Galton (1880) provided the first systematic documentation of such variability, particularly in mental imagery. In recent years, two novel phenomena, hyperphantasia and aphantasia, have become important topics in cognitive neuroscience (Zeman et al., 2015, 2020). In particular, hyperphantasia, characterized by exceptionally vivid mental imagery, might be related to “eidetic imagery,” a major focus of psychological research in the early 20th century (Pearson, 2019; Zeman, 2024).

Eidetic imagery is a form of visual mental imagery that persists in the absence of the stimulus. Unlike afterimages, eidetic images do not move with eye movements and are localized in stably external space (Haber and Haber, 1964; Haber, 1979). The prevalence of eidetic imagery has been reported to be 8% among American elementary school children (Haber and Haber, 1964) and 2–5% among Japanese university students (Matsuoka and Hatakeyama, 2011). Of these, only a very small proportion of individuals with eidetic imagery, less than 1%, possess highly accurate photographic-like images of the original stimulus (Haber, 1979). However, eidetic imagery is considered a constructive process rather than a “photographic” one (Marks, 2023). Eidetic imagery is characterized as a percept-like phenomenon that is “seen” in the literal sense of the word (Jaensch, 1930; Matsuoka and Hatakeyama, 2011), whereas typical images are observed in the mind’s eye (Kosslyn et al., 2006).

Because there are only a few neurophysiological studies on eidetic imagery, the neural basis of eidetic imagery remains unclear. To date, only a single-subject electroencephalogram (EEG) study has reported the occipital alpha wave activity during eidetic imagery (Pollen and Trachtenberg, 1972). In contrast, the neural basis of typical imagery is well-studied. Neuroimaging studies suggest that imagery and perception share partially overlapping neural substrates and have demonstrated that even typical imagery activates the primary visual cortex (Kosslyn et al., 1993; Klein et al., 2004; Slotnick et al., 2005). Moreover, studies using multivoxel pattern analysis (MVPA) have reported the existence of shared neural representations between perception and imagery through cross-decoding approaches (Stokes et al., 2009; Lee et al., 2012; Johnson and Johnson, 2014). These studies have revealed involvement of widespread brain regions, including both the early visual cortex and high-level visual areas. If eidetic imagery possesses perceptual qualities beyond mere subjective impressions, it is hypothesized that its neural substrates share greater commonality with perception than those of typical imagery.

To test this hypothesis, we investigated common neural representations between visual perception and imagery by comparing decoding accuracy in visual areas between eidetikers and typical imagers (controls). Participants performed visual judgements on objects using the stimulus (perception task) or its mental image (imagery task). We first trained the decoder to classify the eight objects in the perception task, and the same classifier was then used to classify the same objects in the imagery task. Such successful cross-decoding indicates the common neural representation of visual perception and imagery, especially in the eidetikers.

## Materials and Methods

### Participants

Participants included 6 eidetikers (3 males and 3 females; mean age, 22.17 years; SD, 1.72) and 13 controls (9 males and 4 females, mean age 21.85 years; SD, 2.3). All participants self-reported as right-handed and were assessed as right-handed using a modified version of the Edinburgh Handedness Inventory (Oldfield, 1971) for Japanese participants (Hatta and Nakatsuka, 1975). There were no significant differences in mean age (Welch’s *t*-test, *t*(12.98) = -0.34, *p* = 0.741) and the handedness score (independent t-test, *t*(17) = 1.69, *p* = 0.110) between the eidetikers and controls. VVIQ (Vividness of Visual Imagery Questionnaire; Marks, 1973) scores (range = 16–80; higher scores indicate more vivid imagery) were 57.83 (SD = 8.33) for the eidetikers and 56.08 (SD = 6.91) for the controls, with no significant difference (Welch’s *t*-test, *t* (8.33)= -0.45, *p* = 0.664). Our participants’ VVIQ scores were moderate compared to those of previous studies (Fulford et al., 2018). Written informed consent was obtained from all participants in accordance with the Declaration of Helsinki. The experimental protocol received approval from the local ethics committee.

### Recruiting of participants

To assess the possession of eidetic imagery, we conducted a preliminary survey that included the Easel Test (Matsuoka et al., 1987). In this survey, we recruited students at Hokkaido University, and 140 participants were assessed to determine whether they possessed eidetic imagery. Among them, 40 who reported “currently experiencing eidetic imagery” or “having experienced it in the past but not at present” in the initial questionnaire were regarded as potential eidetikers, while the remaining 100 who reported “having never experienced it” were regarded as candidates for the controls. The potential eidetikers underwent an interview about their experiences with eidetic imagery to confirm whether they experienced it in daily life, followed by a test battery for eidetic imagery. The control candidates underwent only the test battery. In this battery, 15 of the potential eidetikers were confirmed as eidetikers, and 66 of the control candidates were confirmed as non-eidetikers. Among these confirmed eidetikers and non-eidetikers (controls), only those who consented to participate in the fMRI experiment were included.

The test battery included an afterimage test and the Easel Test. The afterimage test was conducted to help participants understand what a visible image is. In this test, participants gazed at colored panels and then reported what they saw after the panels were removed. The Easel Test was conducted to confirm whether eidetic images were evoked for each participant. The procedure followed Matsuoka et al. (1987), which is a slightly modified version of the original test developed by Haber and Haber (1964). In this test, participants observed picture stimuli for 30 seconds and then verbally reported what they saw in front of their eyes after the stimuli were removed. Participants were confirmed as eidetikers if their reports fulfilled all of the following conditions: the eidetic image did not disappear or move with eye movements, lasted for more than 30 seconds, appeared in positive color, was externally localized, and was reported in the present tense. The eidetic ability of the confirmed eidetikers corresponded to either weak eidetic (i.e., “report a few meaningful weak images”) or strong eidetic (i.e., “report a few or more meaningful vivid images”), as defined by Matsuoka et al. (1987). Participants were confirmed as non-eidetikers if they reported no images. Some participants from both the potential eidetikers and the control candidates reported a few meaningless images (spots, shades, colors, or patterns) and were confirmed as having marginal eidetic imagery (Matsuoka et al., 1987). These participants were excluded from both the eidetikers and controls. The classification of participants was determined by two members of the present study. Verbal reports from a participant classified as an eidetiker are provided in the Supplemental Methods.

### Task procedures

In our main experiment, the eidetikers and controls underwent fMRI scanning, which consisted of 1 resting-state (RS) scanning session, 2 perception task sessions, 6 imagery task sessions, 2 brief retinotopic mapping sessions, a T1 scan, and a DWI scan. Of these, this study only analyzed task scans and T1 scans. The participants performed two tasks: a perception task and an imagery task (Figure 1). They judged whether a red frame presented over a stimulus contained more white or black area. The stimuli included eight monochrome visual objects: apple, melon, pear, banana, bell, wallet, bulb, and phone. These were selected from both natural (fruit) and artificial objects, and half of them were round, while the rest were not.

**Figure 1:**
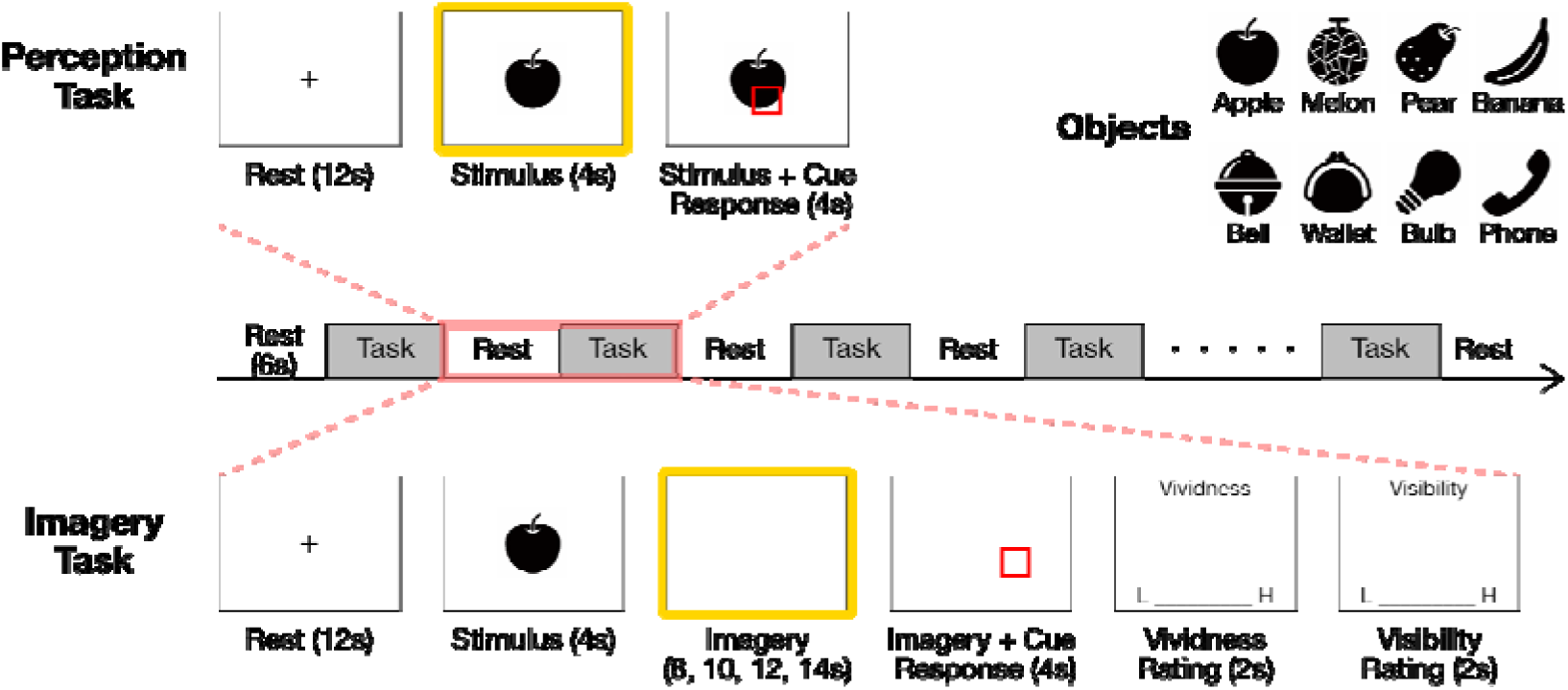
Schematic description of experimental design and time-course. The top-left panel illustrates the perception task, the bottom panel depicts the imagery task, and the top-right panel shows the visual objects used as stimuli in both tasks. The yellow frames indicate the time periods modeled in the general linear model (GLM) in the fMRI analysis.

In the perception task, a trial began with a 12-second rest period, followed by a 4-second stimulus presentation. During the rest period, the participants gazed at the fixation cross, and during the stimulus period, they observed the object with free eye movements. After the 4-second stimulus period, a red frame was presented for 4 seconds somewhere over the stimulus as a response cue. The participants judged the proportion of black and white within the frame based on the actual stimulus, pressing a button with the index finger for black and the middle finger for white. All the stimuli and cues were displayed on an LCD monitor visible via a mirror. Each perception task session consisted of 16 blocks (8 stimuli × 2 trials), and the order of the blocks was randomized within each session. The sessions lasted approximately 12 minutes in total.

In the imagery task, each trial began with a 12-second rest period followed by a 4-second stimulus period. Unlike the perception task, a blank screen was then presented for 8, 10, 12, or 14 seconds to encourage the participants to project and retain an image of the object. This period served as the retention interval, during which participants generated and maintained imagery of the object. After this retention interval, a red frame was presented for 4 seconds somewhere over the area where the object was located as a response cue, while the participants continued to project the image. The eidetikers made their judgments based on what they saw, i.e., an eidetic image, while the controls made their judgments based on a typical mental visual image. At the end of each trial, the participants rated the vividness and visibility of their image within 2 seconds for each rating. The participants responded by pressing a button with their index, middle, ring, and little fingers for each option. For vividness and clarity, they rated how vivid and clear the image was, with the following options: “no image at all,” “vague and dim,” “somewhat clear,” and “clear.” For visibility, they indicated whether they could see the image in front of their eyes rather than in their mind, with the following options: “nothing was visible in front of my eyes,” “it felt like I could see something,” “I could see an image that was irrelevant to the stimulus I was supposed to imagine,” and “I could see the image.” Each imagery task session consisted of 8 blocks (8 stimuli × 1 trial), and the order of the blocks was randomized within each session. The sessions lasted approximately 30 minutes in total. The experiment was conducted consecutively in one day and took approximately 120 minutes to complete, including explanations and all scans.

### MRI acquisition

All scans were performed on a Siemens 3-Tesla Prisma scanner (Erlangen, Germany) with a 64-channel head coil at Hokkaido University. T2*-weighted echo-planar imaging (EPI) was used to acquire a total of 163 scans per perception task session and 143 scans per imagery task session with a gradient EPI sequence. The first three scans within each session were discarded to allow for T1 equilibration. The following scanning parameters were used: repetition time (TR), 2000 ms; echo time (TE), 30 ms; flip angle (FA), 90°; field of view (FOV), 192 × 192 mm; matrix, 94 × 94; 32 axial slices; and slice thickness, 3.0 mm with a 0.88 mm gap. T1-weighted anatomical imaging with an MP-RAGE sequence was performed using the following parameters: TR, 2300 ms; TE, 2.41 ms; FA, 8°; FOV, 233 × 240 mm; matrix, 280 × 288; 224 axial slices; and slice thickness, 0.8 mm without a gap.

### fMRI data processing

Image preprocessing was performed using the SPM12 software (Welcome Department of Cognitive Neurology, http://www.fil.ion.ucl.ac.uk/spm). All functional images were initially realigned to adjust for motion-related artifacts. Volume-based realignment was performed by co-registering images using a rigid body transformation to minimize the squared differences between volumes. The realigned images were then spatially normalized with the Montreal Neurological Institute template based on the affine and nonlinear registration of coregistered T1-weighted anatomical images (normalization procedure of SPM). The images were resampled into 3-mm-cube voxels with the sinc interpolation. They were spatially smoothed using a Gaussian kernel of 6 × 6 × 6 mm full width at half-maximum. In contrast, the images used for MVPA were not smoothed to avoid blurring the fine-grained information contained in the multivoxel activity (Mur et al., 2009; Kamitani and Sawahata, 2010).

Using the general linear model (GLM), the periods indicated by the yellow frames in Figure 1 for each block were modeled as separate box-car regressors that were convolved with a canonical hemodynamic response function. In the imagery task, we added the stimulus period to the model as a covariate of no interest to eliminate the effect of the visual input of the stimulus. Low-frequency noise was removed using a high-pass filter with a cut-off period of 128 s, and serial correlations among scans were estimated using an autoregressive model implemented in SPM12. This analysis yielded 16 (for the perception task session) or 8 (for the imagery task session) independently estimated parameters (beta values) per session for each individual voxel. These parameter estimates were then z-normalized across voxels for each trial and subsequently used as inputs to MVPA.

### fMRI mass-univariate analysis

We used the conventional mass-univariate analysis of individual voxels to identify activated areas for perception and imagery. We analyzed areas that were significantly activated during the stimulus period of the perception task and the imagery period of the imagery task, compared with activation during the rest period. We then directly compared the two tasks. Areas that were significantly activated during the stimulus period of the perception task and the imagery period of the imagery task were compared with activation during the rest period. Contrast images of each participant, generated using a fixed-effects model, were analyzed with a random-effects model using a one-sample *t*-test. Activation for the perception task was reported with a threshold of *p* < .05 corrected for family-wise error (FWE) with an extent threshold of 10 voxels, and for the imagery task and the direct comparison, with a threshold of *p* < .001 uncorrected for multiple comparisons with an extent threshold of 10 voxels.

Additionally, we conducted a region of interest (ROI) analysis to compare brain activity for perception and imagery between the eidetikers and controls. We selected three ROIs from the visual areas, Brodmann areas 17, 18, and 19, using the SPM anatomy toolbox (Eickhoff et al., 2005). Averaged parameter estimates (beta values) for perception and imagery within each ROI were obtained from the results of the fixed-effects model for each participant and compared between the eidetikers and controls using Welch’s *t*-test.

### MVPA

We used MVPA to classify the brain activity during perception and imagery. The classifier was based on a linear support vector machine implemented using LIBSVM (http://www.csie.ntu.edu.tw/~cjlin/libsvm) with a fixed regularization parameter, C = 1. The parameter estimates (beta values) of each trial of voxels within ROIs were used as inputs to the classifier. The ROIs were the same as those used in the mass-univariate analysis.

We tried to classify the eight objects using cross-validation between the perception and imagery task sessions. The classifier was first trained to classify the eight objects in the perception task sessions, and the same classifier was then used to classify the eight objects in the imagery task sessions to test whether the classifier for the object types could generalize between perception and imagery (Figure 2).

**Figure 2:**
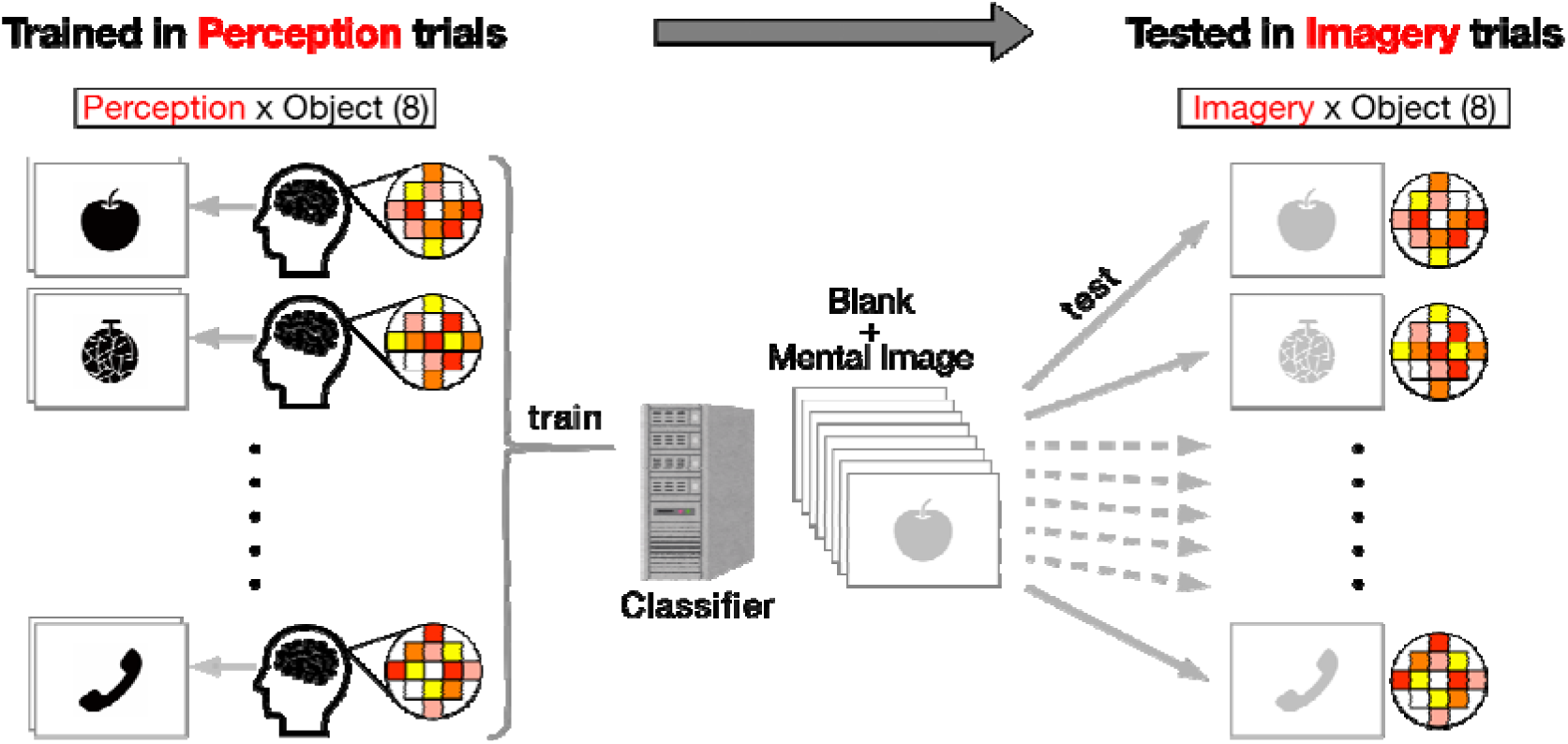
Cross-validation of eight objects between the perception and imagery task sessions. The classifier was trained on the perception task sessions to distinguish the eight objects, and then tested on the imagery task sessions to evaluate its ability to generalize object classification across tasks.

Welch’s *t*-test was used to compare classification accuracy between the eidetikers and controls, with Bonferroni correction applied based on the number of ROIs. For ROIs with a significant group difference, a one-sample *t*-test was used for both groups to determine whether the observed classification accuracy was significantly higher than chance (12.5%), with intersubject differences treated as a random factor (eidetikers, *df* = 5; controls, *df* = 12). Given that eidetic imagery is a type of imagery seen as if it were actual perception, the classification accuracy should be higher for the eidetikers than for the controls.

### Statistical analyses

Behavioral data in the imagery task (i.e., accuracy, response time, vividness and clarity ratings, and visibility ratings) and MRI ROI analyses were conducted using Python (Google Colab, Python 3.12.11). To account for the repeated observations within participants, vividness and clarity ratings were analyzed using linear mixed models (LMM) with restricted maximum likelihood (REML) estimation, whereas visibility ratings were analyzed using cumulative link mixed models (CLMM). These analyses were implemented in R (version 4.5.1) through Python. In both models, group was included as a fixed effect and participants as a random effect. Retention time was not included because its variance was zero in the LMM, and it had too few levels in the CLMM.

## Results

### Behavioral results

To compare the imagery ability between the eidetikers and controls, we analyzed task accuracy and reaction time for correct responses in the imagery task (Figure 3). A two-way ANOVA was conducted with retention time (8, 10, 12, and 14 seconds) and group (eidetikers and controls) as factors. For the task accuracy, there were no significant main effects of retention time (*F*(3, 51) = 0.92, *p* = 0.436) or group (*F*(1, 17) = 0.61, *p* = 0.446), and no significant interaction (*F*(3, 51) = 1.03, *p* = 0.389). For the reaction time, the result showed no significant main effects of retention time (*F*(3, 51) = 0.87, *p* = 0.461) or group (*F*(1, 17) = 1.36, *p* = 0.260), and no significant interaction (*F*(3, 51) = 0.68, *p* = 0.568).

**Figure 3:**
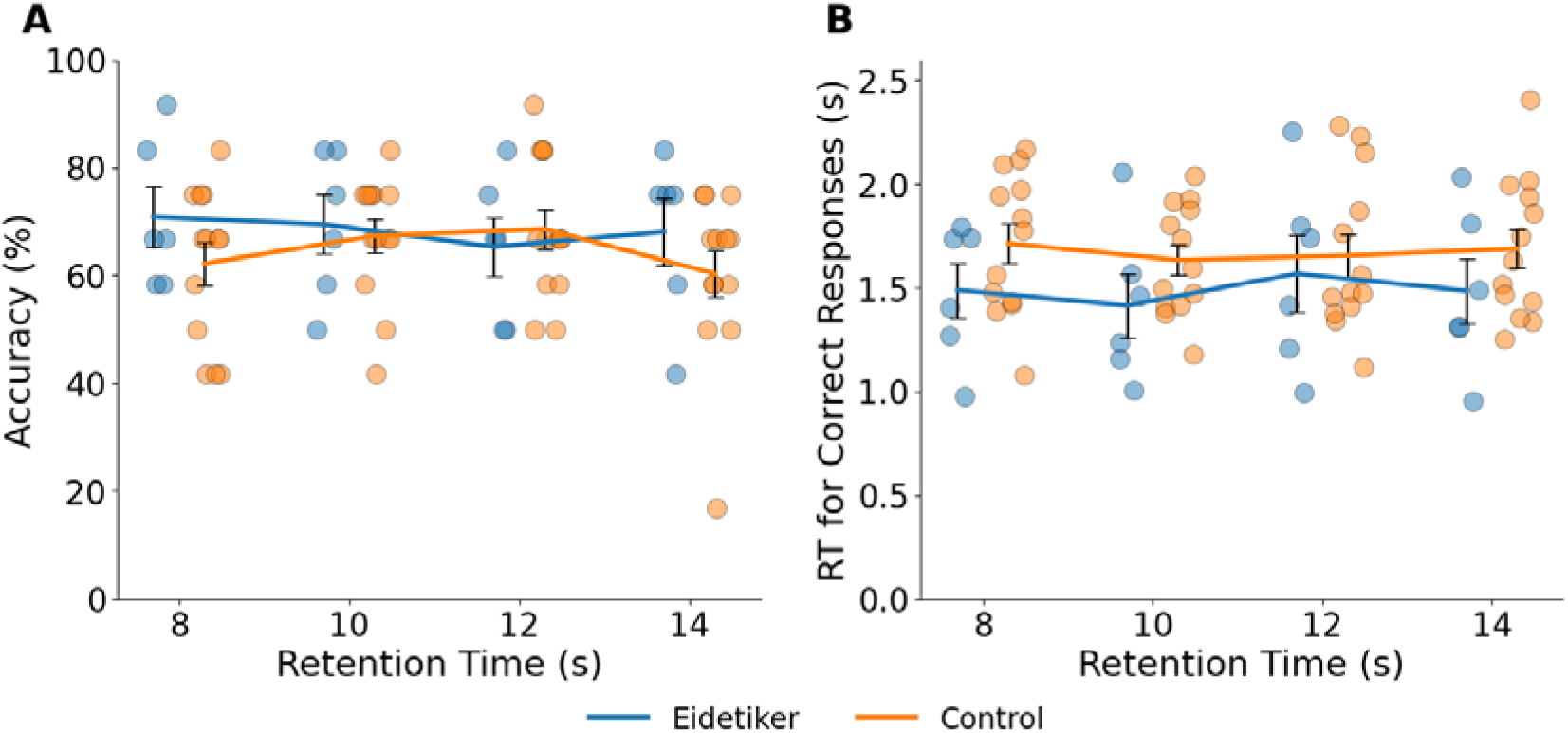
Behavioral Results of the Two Groups. (A) Accuracy during the imagery task for eidetikers (blue line) and controls (orange line). (B) Response times during the imagery task for eidetikers (blue line) and controls (orange line). Each dot represents the result for a single participant. Error bars represent SEM.

The LMM analysis for vividness and clarity ratings (n = 827) revealed no significant effect of group (estimate = 0.58, SE = 0.28, *p* = 0.056). In contrast, the CLMM analysis for visibility ratings (n = 871) showed a significant group effect, with eidetikers having significantly higher odds of giving higher visibility ratings than controls (estimate = 3.44, SE = 0.85, *z* = 4.03, *p* < 0.001). This pattern suggests that eidetikers’ ratings tended to reflect seeing the image in front of their eyes rather than in their mind.

### fMRI mass-univariate analysis

We first analyzed the activated areas for perception and imagery across all participants, compared to the rest period (Tables 1 and 2, Figure 4). For perception, the activated areas included the bilateral fusiform gyrus, bilateral middle occipital gyrus, and right lingual gyrus. For imagery, no significant activity was observed using the same threshold as for perception. However, using a more liberal threshold, significant activity was found in the bilateral lingual gyrus, right cuneus, bilateral superior occipital gyrus, and left superior frontal gyrus.

**Table 1:**
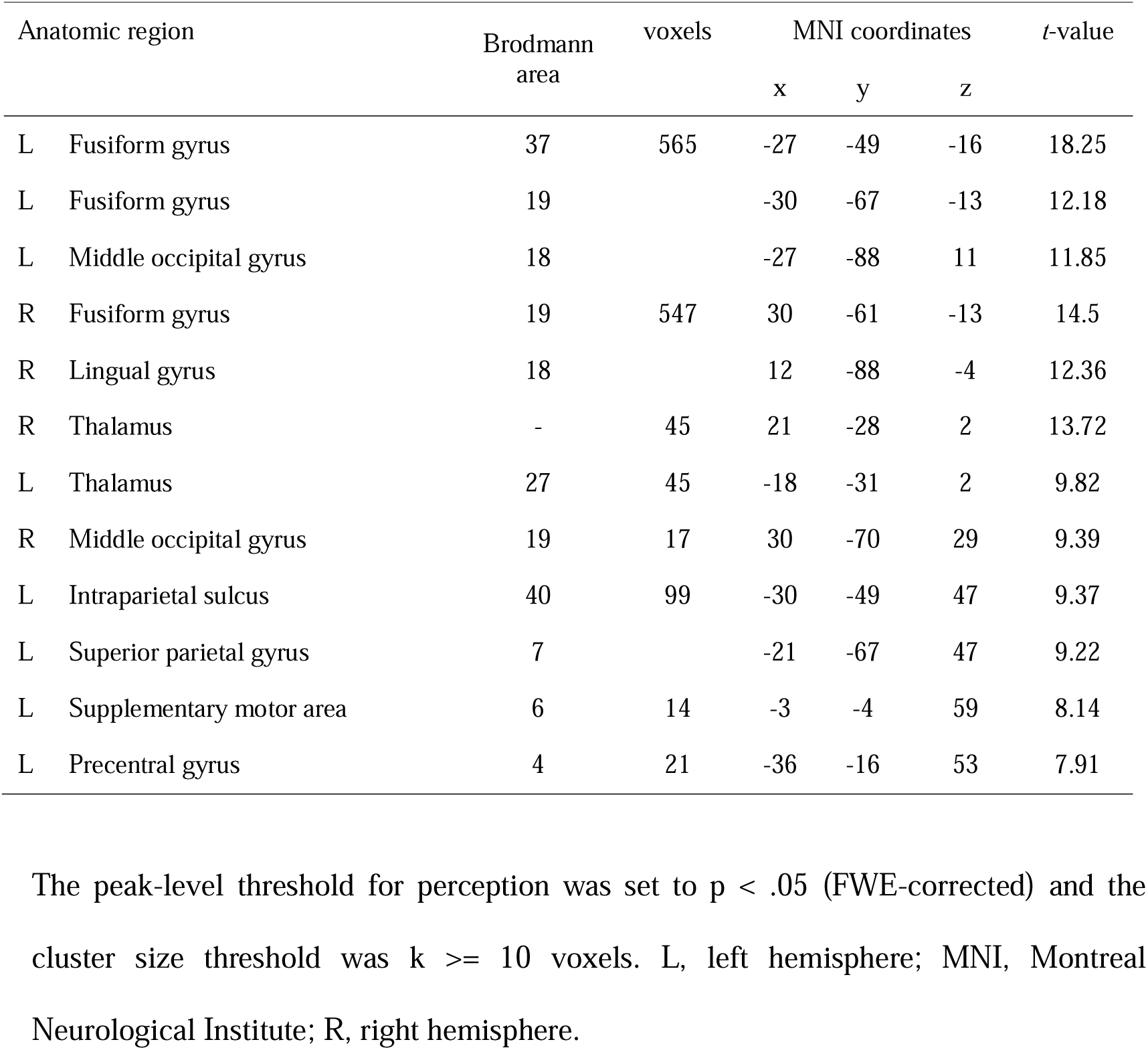
Anatomical regions, peak voxel coordinates, and t-values of observed activations during the perception task across all participants

| Anatomic region |  | Brodmann area | voxels | MNI coordinates |  |  | t-value |
| --- | --- | --- | --- | --- | --- | --- | --- |
|  |  |  |  | x | y | z |  |
| L | Fusiform gyrus | 37 | 565 | -27 | -49 | -16 | 18.25 |
| L | Fusiform gyrus | 19 |  | -30 | -67 | -13 | 12.18 |
| L | Middle occipital gyrus | 18 |  | -27 | -88 | 11 | 11.85 |
| R | Fusiform gyrus | 19 | 547 | 30 | -61 | -13 | 14.5 |
| R | Lingual gyrus | 18 |  | 12 | -88 | -4 | 12.36 |
| R | Thalamus | - | 45 | 21 | -28 | 2 | 13.72 |
| L | Thalamus | 27 | 45 | -18 | -31 | 2 | 9.82 |
| R | Middle occipital gyrus | 19 | 17 | 30 | -70 | 29 | 9.39 |
| L | Intraparietal sulcus | 40 | 99 | -30 | -49 | 47 | 9.37 |
| L | Superior parietal gyrus | 7 |  | -21 | -67 | 47 | 9.22 |
| L | Supplementary motor area | 6 | 14 | -3 | -4 | 59 | 8.14 |
| L | Precentral gyrus | 4 | 21 | -36 | -16 | 53 | 7.91 |

The peak-level threshold for perception was set to $p < .05$ (FWE-corrected) and the
cluster size threshold was $k \geq 10$ voxels. L, left hemisphere; MNI, Montreal
Neurological Institute; R, right hemisphere.

**Table 2:** Anatomical regions, peak voxel coordinates, and t-values of observed activations during the imagery task across all participants

| Anatomic region |  | Brodmann area | voxels | MNI coordinates |  |  | *t*-value |
| --- | --- | --- | --- | --- | --- | --- | --- |
|  |  |  |  | x | y | z |  |
| R | Lingual gyrus | 18 | 54 | 15 | -76 | -1 | 6.68 |
| R | Cuneus | 18 | 76 | 15 | -91 | 14 | 6.3 |
| R | Superior occipital gyrus | 18 |  | 21 | -88 | 29 | 4.82 |
| L | Lingual gyrus | 18 | 26 | -15 | -76 | -4 | 6.21 |
| L | Superior occipital gyrus | 17 | 39 | -9 | -91 | 8 | 5.87 |
| L | Superior frontal gyrus, dorsolateral | 6 | 35 | -24 | -4 | 59 | 5.43 |
| R | Superior parietal gyrus | 5 | 24 | 21 | -58 | 59 | 4.73 |
| L | Middle occipital gyrus | 19 | 15 | -21 | -85 | 20 | 4.63 |
| R | Middle temporal gyrus | 37 | 20 | 45 | -70 | 11 | 4.07 |

The peak-level threshold for imagery was set to $p < .001$ (uncorrected) and the cluster size threshold was $k \geq 10$ voxels. L, left hemisphere; MNI, Montreal Neurological Institute; R, right hemisphere.

**Figure 4:**
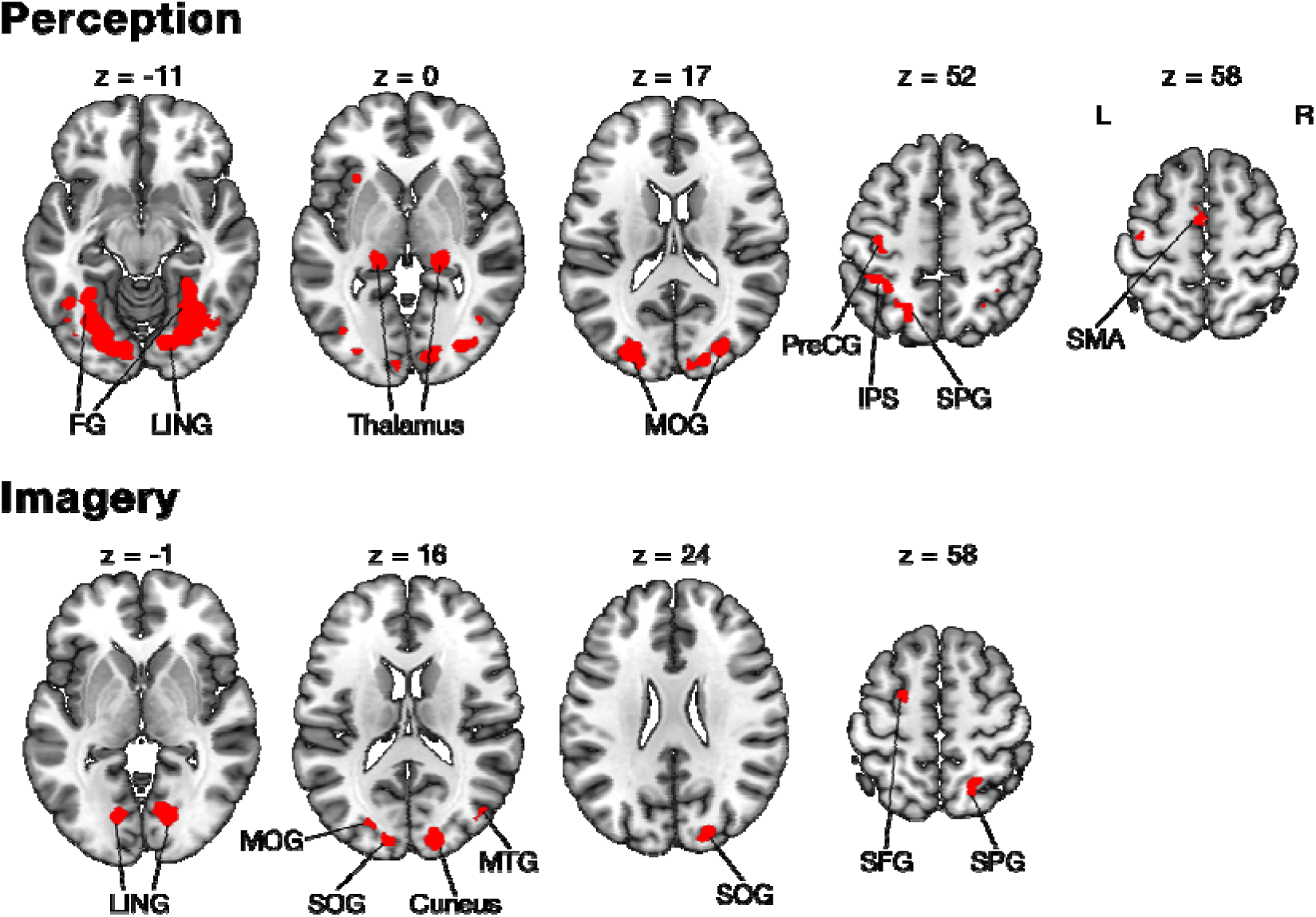
Activated areas for the perception and imagery tasks. FG, Fusiform gyrus; LING, Lingual gyrus; MOG, Middle occipital gyrus; PreCG, Precentral gyrus; IPS, Intraparietal sulcus; SPG, Superior parietal gyrus; SMA, Supplementary motor area; SOG, Superior occipital gyrus; MTG, Middle temporal gyrus; SFG, Superior frontal gyrus

We then directly compared brain activity between perception and imagery, generating two contrast images for all participants: perception versus imagery and imagery versus perception. However, both images showed no significant difference in the activated areas between perception and imagery.

Additionally, we compared the parameter estimates (beta values) within the four ROIs for the eidetikers and controls, for both perception and imagery (Figure 5A). The average beta values for each ROI showed no significant differences between eidetikers and controls for either perception or imagery (*ps* > 0.05).

**Figure 5:**
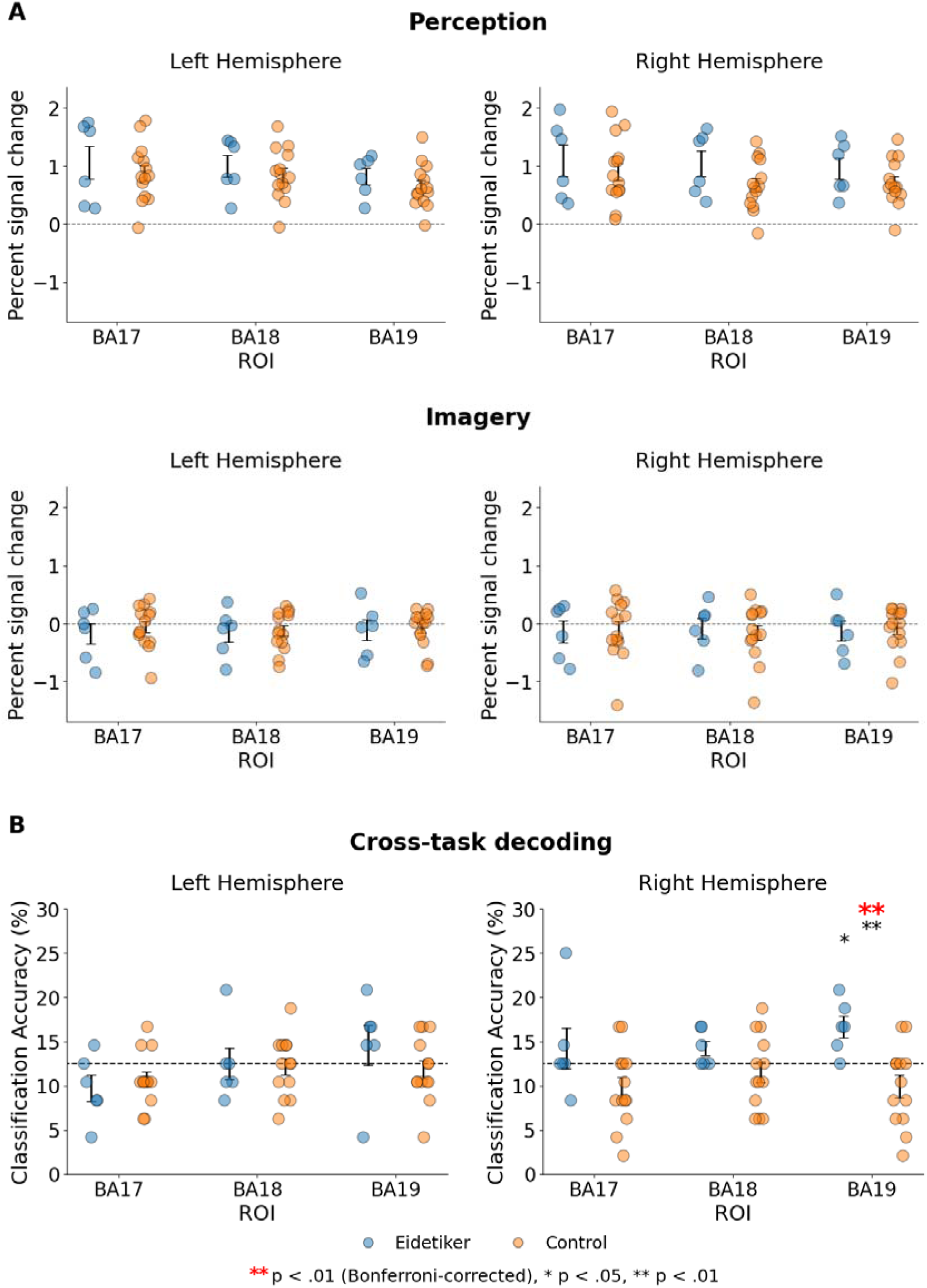
fMRI Percent Signal Change and Decoding Results. A) Percentages of BOLD signal changes in eidetikers and controls during perception and imagery tasks. B) Results of cross-task decoding between imagery and perception processes. Red and black asterisks indicate statistically significant results after Bonferroni correction and uncorrected significance, respectively. Significance levels are indicated as *p < 0.05, **p < 0.01. The dashed lines indicate the chance level. Blue and orange dots represent individual data points from eidetikers and controls, respectively. Error bars represent SEM.

### MVPA

An ROI-based MVPA was conducted to classify the eight objects across perception and imagery. We expected the classification accuracy to be higher for eidetikers than for controls, given that eidetic imagery is a type of imagery experienced as if it were actual perception. Figure 5B shows the classification accuracies for each ROI for eidetikers and controls. Significant differences between the eidetikers and controls were found in the right BA19 (Welch’s t-test, *t*(15.12) = 3.88, *p* = 0.009, Bonferroni-corrected), with the eidetikers’ accuracy higher than that of the controls in the ROIs. Notably, the eidetikers’ accuracy in BA19 was significantly above chance (one-sample t-test, *t*(5) = 3.46, *p* = 0.018), whereas the controls’ accuracy did not differ significantly from chance (one-sample t-test, *t*(12) = -2.05, *p* = 0.063).

## Discussion

This study used fMRI with MVPA to test whether neural representations of visual perception and imagery are more strongly shared in individuals with eidetic imagery than in controls. Our results showed that imagined objects could be successfully classified based on activation patterns during perception in the right BA19 for eidetikers, but not for controls. This result was consistent with the significant group difference in the visibility rate, which reflected the degree to which participants reported that the image appeared to be “seen in front of their eyes.” That is, the subjective percept-like experience of images—such as eidetic imagery—suggests a higher degree of similarity between neural representations of perception and imagery.

Previous MVPA studies on typical imagery have shown similar neural representations between perception and imagery across widespread visual areas, with relatively greater overlap in high-level visual areas (Lee et al., 2012; Dijkstra et al., 2019). Among these regions, the lateral occipital complex (LOC) has been repeatedly shown to contain perceptual activation patterns that can be used to decode the content of imagery (Stokes et al., 2009; Reddy et al., 2010; Ragni et al., 2020). In the present study, the BA19 identified in eidetikers closely corresponds to the LOC (Malach et al., 1995). This is consistent with the current task that required imagery of object shapes. In contrast, our controls with typical imagery did not show successful decoding from perception to imagery, unlike eidetikers. Two possible reasons may account for this result. First, in our experimental design, the time course differed between the perception and imagery tasks, with imagery trials varying in duration and lasting longer than perception trials. Second, in analyzing the imagery task, we added the stimulus period to the fMRI model as a covariate of no interest to eliminate the effect of visual input from the stimulus. This modeling procedure may attenuate the effective signal available for cross-decoding. Despite these reasons, successful cross-decoding in eidetikers may reflect a high degree of shared neural representations between perception and imagery.

Interestingly, the current cross-decoding analysis revealed a group difference between eidetikers and controls in BA19, whereas no such difference was found in the early visual cortex, including BA17 and BA18. In comparison, Lee et al. (2012) reported that cross-decoding between perception and imagery correlated with the vividness of visual imagery, particularly in the early visual cortex. This difference may indicate that eidetic imagery cannot be regarded as solely an extreme case on the vividness dimension. In typical imagery, the early visual cortex is recruited particularly when the imagery requires fine-grained visual details (Ragni et al., 2020; Dijkstra, 2024). Our eidetikers were not limited to those having highly detailed, photographic-like images, because the Easel Test used in our study did not require images to be particularly detailed. Therefore, it remains possible that if we examine only individuals with more finely detailed eidetic imagery, group differences might also emerge in the early visual cortex. Importantly, although the involvement of early visual areas in imagery vividness appears to be variable, relatively high-level visual areas, such as BA19, are involved more consistently (Fulford et al., 2018). This may relate to the fact that imagery vividness is influenced not only by the amount of visual information contained in the representation, such as clarity and colourfulness, but also by other factors such as liveliness (Marks, 2023). Taken together, these findings suggest that the observed BA19 differences may reflect individual variability in subjective imagery experience, including aspects such as liveliness, rather than merely representing the extreme of image detail.

A limitation of the present study is that we relied on anatomical ROIs, which did not permit further distinction of functional subregions within BA19. Achieving this level of specificity requires additional functional localizer tasks beyond standard retinotopic mapping (Engel et al., 1997; Tootell et al., 1998). Such functional localizer tasks could reveal more localized group differences within BA19, potentially identifying regions involved in the emergence of eidetic imagery as well as those underlying individual differences in subjective imagery experience.

In summary, our results reveal a similarity between the neural representations of visual perception and imagery in eidetikers, providing neuroscientific evidence for the existence of eidetic imagery. Although group differences between eidetikers and controls were not apparent in standard measures such as behavioral data or average brain activation, this may reflect the elusive nature of eidetic imagery, historically described as the “will-o’-the-wisp” (Doob, 1966) or the “ghost” (Haber, 1979). In the present study, group differences were detected using MVPA and subjective visibility ratings, highlighting the neural basis of eidetic imagery that extends beyond mere phenomenological description.

## Acknowledgments

This work was supported by JSPS KAKENHI Grant Number JP23H00078 to K.O. We thank Drs. Kazuo Matsuoka, Kohsuke Takahashi, and Fumihito Imai for helpful comments during the conduct of this study. The authors used AI-assisted language editing tools (ChatGPT and DeepL) to improve the clarity and readability of the manuscript.

## References

Dijkstra N (2024) Uncovering the role of the early visual cortex in visual mental imagery. Vision (Basel) 8:29.

Dijkstra N, Bosch SE, van Gerven MAJ (2019) Shared neural mechanisms of visual perception and imagery. Trends Cogn Sci 23:423–434.

Doob LW (1966) Eidetic imagery: a cross-cultural will-o’-the-wisp? J Psychol 63:13–34.

Eickhoff SB, Stephan KE, Mohlberg H, Grefkes C, Fink GR, Amunts K, Zilles K (2005) A new SPM toolbox for combining probabilistic cytoarchitectonic maps and functional imaging data. NeuroImage 25:1325–1335.

Engel SA, Glover GH, Wandell BA (1997) Retinotopic organization in human visual cortex and the spatial precision of functional MRI. Cereb Cortex 7:181–192.

Fulford J, Milton F, Salas D, Smith A, Simler A, Winlove C, Zeman A (2018) The neural correlates of visual imagery vividness - An fMRI study and literature review. Cortex 105:26–40.

Galton F (1880) Statistics of mental imagery. Mind 5:301–318.

Haber R (1979) Twenty years of haunting eidetic imagery: where’s the ghost? Behav Brain Sci 2:583–594.

Haber RN, Haber RB (1964) Eidetic imagery. I. Frequency. Percept Mot Skills 19:131–138.

Hatta T, Nakatsuka Z (1975) Handedness inventory. In: Papers on Celebrating the 63rd Birthday of Prof. Ohnishi (Ohno S, ed), pp 224–247. Osaka: Osaka City University.

Jaensch ER (1930) Eidetic Imagery and Typological Methods of Investigation: Their Importance for the Psychology of Childhood, the Theory of Education, General Psychology, and the Psychophysiology of Human Personality. Transl. from the 2. Ed. by Oscar Oeser.

Johnson MR, Johnson MK (2014) Decoding individual natural scene representations during perception and imagery. Front Hum Neurosci 8:59.

Kamitani Y, Sawahata Y (2010) Spatial smoothing hurts localization but not information: pitfalls for brain mappers. NeuroImage 49:1949–1952.

Klein I, Dubois J, Mangin JF, Kherif F, Flandin G, Poline JB, Denis M, Kosslyn SM, Le Bihan D (2004) Retinotopic organization of visual mental images as revealed by functional magnetic resonance imaging. Brain Res Cogn Brain Res 22:26–31.

Kosslyn SM, Alpert NM, Thompson WL, Maljkovic V, Weise SB, Chabris CF, Hamilton SE, Rauch SL, Buonanno FS (1993) Visual mental imagery activates topographically organized visual cortex: PET investigations. J Cogn Neurosci 5:263–287.

Kosslyn SM, Thompson WL, Ganis G (2006) The case for mental imagery. Oxford University Press.

Lee S-H, Kravitz DJ, Baker CI (2012) Disentangling visual imagery and perception of real-world objects. NeuroImage 59:4064–4073.

Malach R, Reppas JB, Benson RR, Kwong KK, Jiang H, Kennedy WA, Ledden PJ, Brady TJ, Rosen BR, Tootell RB (1995) Object-related activity revealed by functional magnetic resonance imaging in human occipital cortex. Proc Natl Acad Sci U S A 92:8135–8139.

Marks DF (1973) Visual imagery differences in the recall of pictures. Br J Psychol 64:17–24.

Marks DF (2023) Phenomenological studies of visual mental imagery: A review and synthesis of historical datasets. Vision (Basel) 7:67.

Matsuoka K, Hatakeyama T (2011) Research on eidetic imagery in Japan. J Ment Imagery 35:123–136.

Matsuoka K, Onizawa T, Hatakeyama T, Yamaguchi H (1987) Incidence of young adult eidetikers, and two kinds of eidetic imagery. Tohoku Psychologica Folia 46:62–74.

Mur M, Bandettini PA, Kriegeskorte N (2009) Revealing representational content with pattern-information fMRI - An introductory guide. Soc Cogn Affect Neurosci 4:101–109.

Oldfield RC (1971) The assessment and analysis of handedness: the Edinburgh inventory. Neuropsychologia 9:97–113.

Pearson J (2019) The human imagination: the cognitive neuroscience of visual mental imagery. Nat Rev Neurosci 20:624–634.

Pollen D, Trachtenberg M (1972) Alpha rhythm and eye movements in eidetic imagery. Nature 237:109–112.

Ragni F, Tucciarelli R, Andersson P, Lingnau A (2020) Decoding stimulus identity in occipital, parietal and inferotemporal cortices during visual mental imagery. Cortex 127:371–387.

Reddy L, Tsuchiya N, Serre T (2010) Reading the mind’s eye: decoding category information during mental imagery. NeuroImage 50:818–825.

Slotnick SD, Thompson WL, Kosslyn SM (2005) Visual mental imagery induces retinotopically organized activation of early visual areas. Cereb Cortex 15:1570–1583.

Stokes M, Thompson R, Cusack R, Duncan J (2009) Top-down activation of shape-specific population codes in visual cortex during mental imagery. J Neurosci 29:1565–1572.

Tootell RB, Hadjikhani NK, Vanduffel W, Liu AK, Mendola JD, Sereno MI, Dale AM (1998) Functional analysis of primary visual cortex (V1) in humans. Proc Natl Acad Sci U S A 95:811–817.

Zeman A (2024) Aphantasia and hyperphantasia: exploring imagery vividness extremes. Trends Cogn Sci 28:467–480.

Zeman A, Dewar M, Della Sala S (2015) Lives without imagery - Congenital aphantasia. Cortex 73:378–380.

Zeman A, Milton F, Della Sala S, Dewar M, Frayling T, Gaddum J, Hattersley A, Heuerman-Williamson B, Jones K, MacKisack M, Winlove C (2020) Phantasia - The psychological significance of lifelong visual imagery vividness extremes. Cortex 130:426–440.

