## Supplemental Methods for "Common neural representation between visual perception and imagery in eidetikers"

The following verbatim reports were obtained from the same participant in response to two different stimuli.
The Easel Test was conducted in Japanese, and the verbatim transcripts were translated into English using a generative AI system, with minimal editing to preserve the original content.

**<The silhouette picture of a Crocodile and a Boy>**

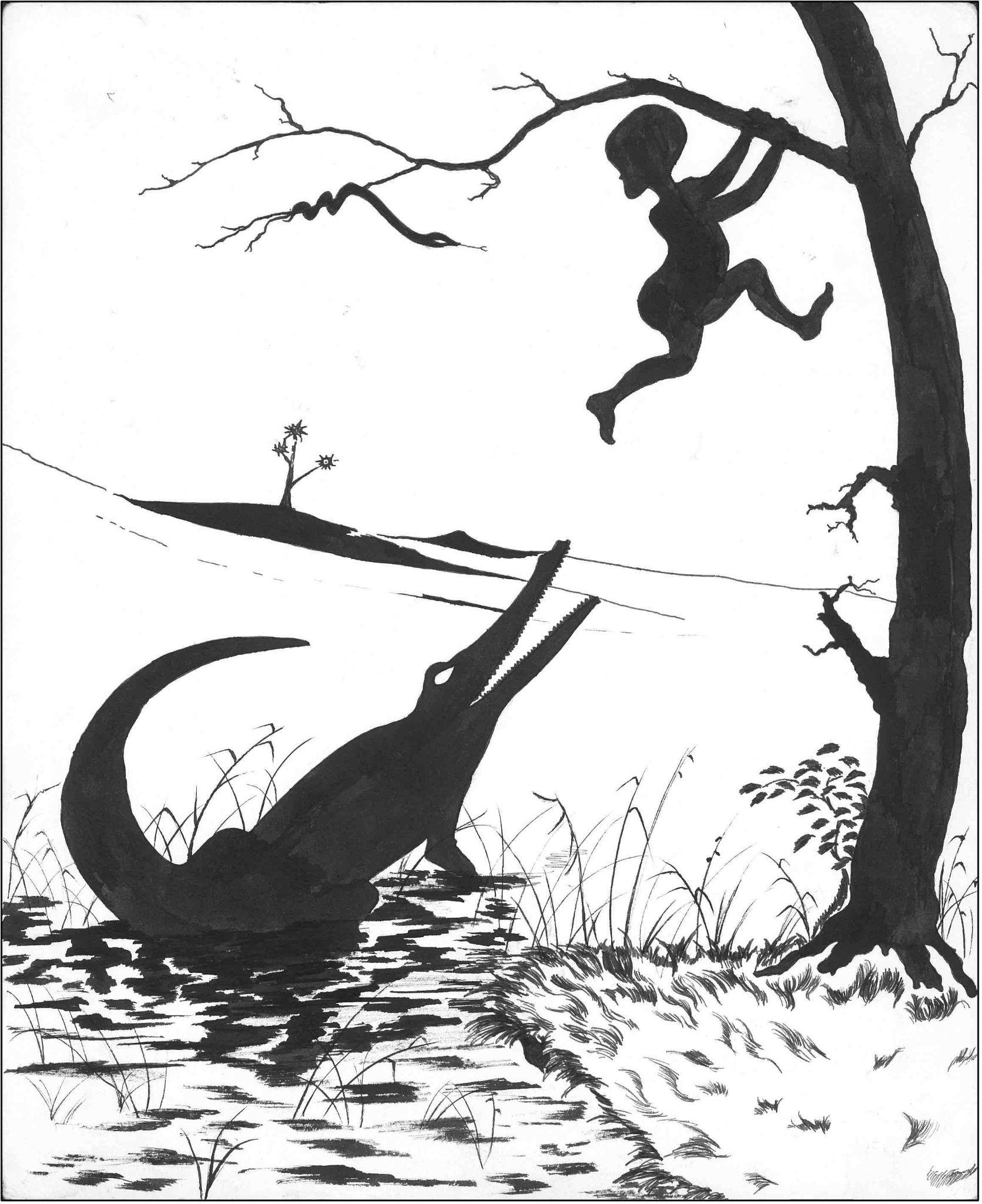
(Kluver 1926)

(After the afterimage test)

**Experimenter (E):** Now I’d like you to look at a picture. But this time, unlike earlier where you fixated on one point, please move your eyes and look over the whole picture carefully. And when I remove the picture, just like before, please keep looking at the place where the picture was. At that time, even if you move your eyes within the screen, that’s fine. And after it’s been removed, if you see anything, please describe in as much detail as you can whatever you see. If you don’t see anything in particular, please report that you don’t see anything.

**Participant (P):** Okay.

**E:** Do you have any questions?

**P:** No, I’m good.

**E:** Alright. I’ll show you the picture now. Please move your eyes and look over everything carefully. Here we go.

(After 30 seconds, E removed the stimulus picture.)

**P:** Ah, I can still see the afterimage from earlier.

**E:** What exactly do you see?

**P:** Like… a snake and a crocodile. And, well, kind of like a desert. I guess.

**E:** What color do the snake and crocodile look?

**P:** The snake and crocodile… uh, purple.

**E:** Purple.

**P:** And I can, like, sorta see a person too, but… um, maybe black.

**E:** When you move your eyes, does it follow your eyes? Or does it stay in one place?

**P:** It stays.

**E:** It stays. Are you still seeing it now?

**P:** I can see it faintly… but not as clearly as at first.

**E:** Then could you tell me what you’re seeing right now?

**P:** Right now… the faint crocodile and snake from earlier. But the person is not really… yeah, I don’t really see the person. Right. I can almost no longer see it.

**E:** Is it still very faint there?

**P:** No, I think it’s fine to say I don’t see it anymore. Yeah.

（2’09’’ from the beginning of the report）

**E:** If you try to see the picture from earlier again in front of you, does it appear?

**P:** It does.

**E:** Then could you describe it in detail?

**P:** You mean like what I saw at the very beginning, right?

**E:** Yes. If the things that appeared right after the removal appear again in front of you, please describe them.

**P:** There’s a crocodile, a snake, a person. There’s vegetation around the crocodile. The snake and the person are connected somehow. The person is climbing a tree. In the back there’s an island. But it’s like… more desert-like than sea-like. Yeah.

**E:** What colors do they look like?

**P:** Well, basically purple or black or white or light blue. Like shades of light colors.

**E:** Are the outlines clear? Or do they look blurred?

**P:** When I first remembered it—right after you removed the picture—it was pretty blurry. But when I recall it myself, it looks rather clear.

**E:** Are you still seeing it the same way now? Or is it gradually changing?

**P:** Well… no, I don’t see it much anymore. It’s fainter than when I first recalled it.

**E:** It’s getting faint? Then you can stop looking at it.

**P:** Okay.

**E:** When I say that, does it disappear?

**P:** Yes. I don’t see it. But still… yeah, the crocodile keeps appearing quite a lot. Maybe it left a strong impression.

**E:** The crocodile appears—

**P:** The crocodile and the snake appear clearly.

**E:** Alright, thank you.

（4’23’’ from the beginning of the initial report）

**<The picture of Mice>**

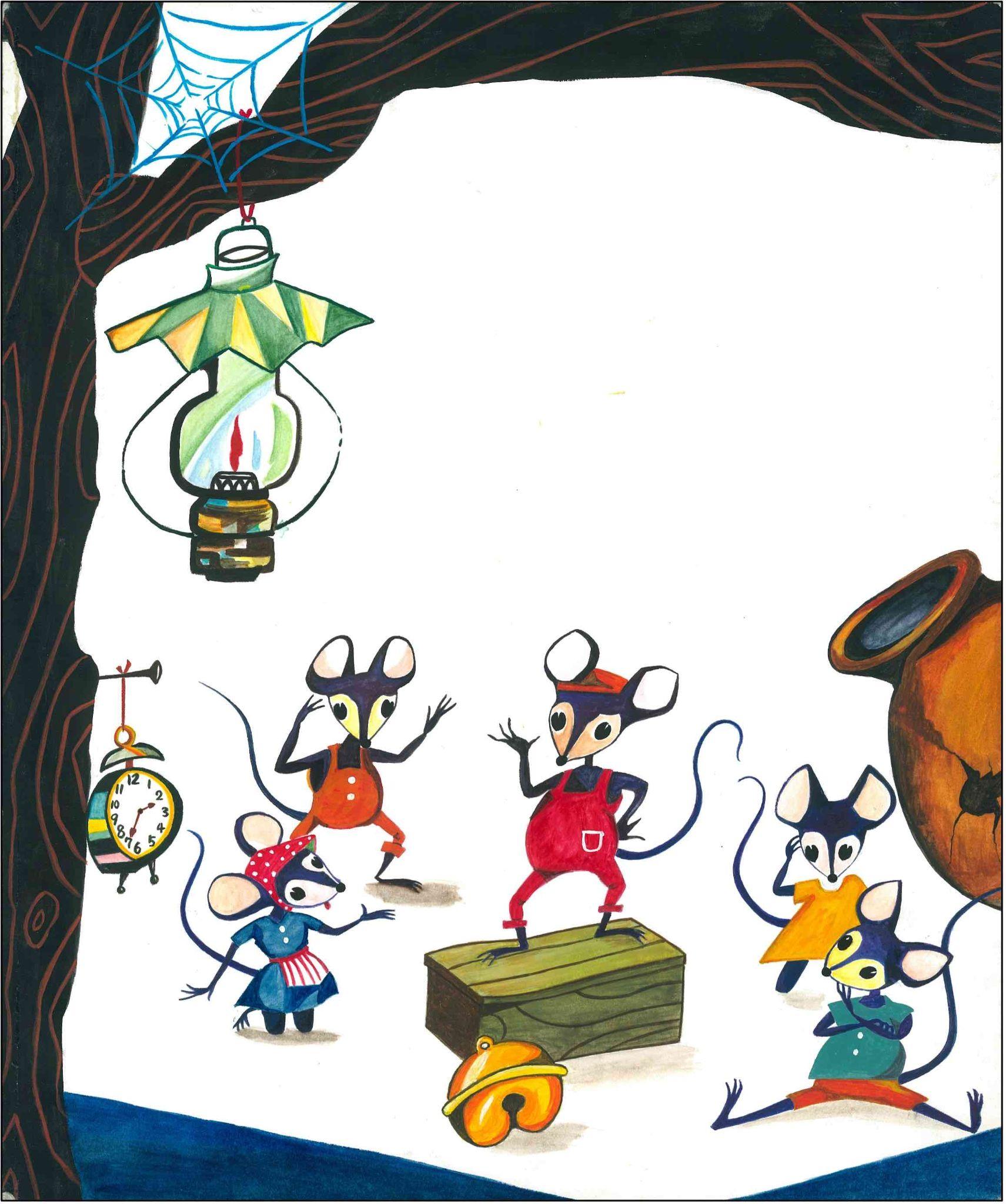
(Onizawa, T., Eto, Y., & Matsuoka, K., 1983)

(E removed the stimulus picture.)

P: Well… yeah, compared to earlier, I can see it more clearly.
There are five mice. And there’s one mouse in the center, and that one is the biggest—well, maybe the most important one. It’s standing on a platform. And under the platform there’s a bell, and there’s a tree. And on the tree there’s something like a clock hanging. And below that there’s something like a curtain. Yeah… that’s about it, I guess.

E: Could you tell me the colors of each of them?

P: The colors? The mouse in the center is red. And the ones next to it are orange… I guess orange? Orange and yellow? But this one over here also looks kind of orange and yellow.
And the curtain is probably blue, and the clock is colorful? And the clock hands point to the numbers 2 and 7. And there’s also a lamp above the clock, and the lamp is relatively bigger than the clock. And yeah, green and yellow leave a strong impression, and the surrounding trees are kind of dark. And instead of straight vertical lines, the lines look more like spiral shapes. And there’s also a spider web, I think. But there’s no spider. That’s how it seems.

E: What color does the spider web look?

P: Light blue? White? Light blue or white.

E: How many layers does it look like?

P: Umm… maybe it’s double-layered. But rather than being double, it’s more like a single thick line—each line feels heavy, like a bold line.

E: Since it started appearing, did anything change? Did it gradually fade overall, or did some parts disappear?

P: Well, overall it faded. Yeah.
But the mice—uh, well—the impression of the one in the center was strong, but the others, not so much. I don’t think I could really tell their expressions… or rather, I don’t really know. Yeah. Yeah, but now I don’t see it anymore.

E: Has everything disappeared now?

P: Yeah.

（3’00’’ from the beginning of the report）
